# Design and Optimization of Multifunctional Peptide Candidates for Cosmeceutical Applications: Combining Anti-Inflammatory and Collagen-Boosting Properties

**DOI:** 10.1101/2025.06.26.661667

**Authors:** Aishwarya Natarajan, Aswini Javvadi, Sarath Kolli

**Affiliations:** Boltzmann Labs, Hyderabad 500084, Telangana, India; Department of Biosciences and Bioengineering, Indian Institute of Technology Bombay (IITB), Powai, Mumbai, Maharashtra 400076, India

## Abstract

Skin aging is multifactorial and involves inflammation, oxidative stress, collagen degradation, and impaired metal ion homeostasis, which requires holistic approaches for dermal rejuvenation. Conventional cosmeceutical peptides often target singular pathways, limiting their efficacy. In this study, we present a computational strategy for designing and evaluating multifunctional peptides that can simultaneously modulate collagen remodelling, exhibit anti-inflammatory responses, and maintain extracellular matrix (ECM) stability. We focused on three important skin-associated proteins: TGF-β receptor II, Integrin α5β1, and MMP-1, and generated peptide sequences specific to them through a de novo sequence generation method that utilized protein-specific profiles alongside machine learning-based property screening. Over 15,000 peptides were generated and filtered using physicochemical and synthesis-based criteria (charge, hydrophobicity, pI, solubility, and synthesis rules). A multi-parametric composite score was developed to visualize peptide favourability in a 3D physicochemical space. The top peptides from each target set were subjected to structure prediction and docking, resulting in binding affinities that ranged from –19.8 to –34.5 kcal/mol. Cross-target docking analysis demonstrated a significant potential for dual or pan-target applications, indicating the broad-spectrum utility of the cosmeceutical. This study firmly establishes a foundation for multifunctional peptide candidates that combine collagen-boosting, anti-inflammatory, and MMP-inhibitory effects. These candidates are poised to lead the way for innovative skincare formulations, paving the path for future validation and experimental evidence that will undeniably strengthen our findings.

## Introduction

Skin aging is a complex, multi-factorial biological process affected by intrinsic genetic factors and extrinsic environmental influences, including UV radiation, oxidative stress, and inflammation. (1-3) Visible signs of aging, including wrinkles, decreased elasticity, and uneven texture, stem from the gradual breakdown of the extracellular matrix (ECM), especially collagen. (4,5) These structural changes result from decreased fibroblast signaling, excessive activation of matrix metalloproteinases (MMPs), and compromised cell and extracellular matrix (ECM) interactions. (6,7) Targeting interconnected pathways offers a strategic approach to reversing or delaying visible signs of aging. Bioactive peptides have emerged as a promising class of cosmeceuticals capable of targeting specific pathways involved in dermal aging. (8-10) Current commercial peptides typically focus on a single action mechanism— collagen stimulation, anti-inflammation, or MMP inhibition. (8,11,12) The single-target approach in dermatological research fails to address the intricate biology of the skin, where signaling pathways interact in complex ways that influence various physiological processes. Consequently, the efficacy of peptide-based therapies is often limited, as they do not adequately consider the broader biological context. (13,14) Furthermore, while several peptides have been reported to bind MMPs or collagen receptors in isolation (15,17), there remains a lack of multifunctional peptide candidates that can simultaneously modulate pro-regenerative (e.g., TGF-β signaling) (16,18) and anti-degradative (e.g., MMP suppression) axes.

Despite recent advances, most peptide discovery efforts still depend primarily on in vitro or empirical screening methods, which often involve expensive and labour-intensive assays. There is a significant deficiency in workflows that systematically utilize rational in silico design and prioritization. This approach should incorporate structural compatibility and physicochemical profiling to identify and select the top candidates efficiently. (19-21) Although individual computational tools like structure-based affinity scoring and target-specific peptide docking have been developed, a cohesive framework that effectively combines de novo sequence generation, multi-objective optimization, and structural docking— particularly for skin-aging applications—remains absent. (22,23) Moreover, there is minimal exploration of how peptide physicochemical properties (e.g., net charge, hydrophobicity, isoelectric point) relate to multifunctionality and target binding, which are critical determinants of skin permeability, stability, and activity. (24,25)

In this study, we address these gaps by presenting a computational pipeline for the rational design and screening of multifunctional peptides targeting three critical skin-associated proteins:

- TGF-β receptor II (TGFBR2): regulator of collagen biosynthesis,
- Integrin α5β1: modulator of fibroblast-ECM signaling,
- MMP-1: a collagenase responsible for ECM degradation.

We identify peptides with dual or triple activity profiles using de novo sequence generation, machine learning–based property filtering, and molecular docking. We further construct a composite score to visualize and rank peptides across a multidimensional physicochemical space, facilitating the selection of top candidates with optimized binding potential and cosmetic compatibility.

This integrative approach offers a new direction for the development of next-generation cosmeceutical peptides—short, stable, multifunctional sequences that may deliver enhanced dermal regeneration through synergistic biological modulation.

## Methods

### 1. Target Selection and Structural Profiling

Three key dermal targets involved in skin aging were selected for peptide design and screening: Transforming Growth Factor-β Receptor II (TGFBR2), Integrin α5β1, and Matrix Metalloproteinase-1 (MMP-1). These were chosen based on their critical roles in collagen synthesis, ECM remodelling, and inflammatory regulation. The crystallographic structures were obtained from the Protein Data Bank (PDB): TGFBR2 (PDB ID: 5E8X); Integrin α5β1 (PDB ID: 3VI4); MMP-1 (PDB ID: 1SU3). Each structure was prepared using standard preprocessing steps, including removal of heteroatoms, addition of hydrogens, and assignment of correct protonation states using AutoDockTools.(26)

### 2. Peptide Sequence Generation

A total of ~15,000 short peptides (10–15 amino acids) were generated through a combination of: Motif-guided assembly: Based on known bioactive skin peptides and receptor-binding motifs. Randomized de novo design: Constrained by biological amino acid frequencies and receptor compatibility heuristics. Sequences were generated using in-house Python scripts and BoltPro’s peptide library module called Peprclip. (27) Custom constraints were introduced to favor residues known to influence skin permeability (e.g., Arg, Pro, Gly) and structural stability.

### 3. Physicochemical Filtering and Composite Scoring

The raw peptide library was filtered based on key properties: Net charge (at pH 7.0); Average hydrophobicity (GRAVY score); Isoelectric point (pI); Solubility prediction (NetSolP); Synthetic feasibility (removal of sequences with multiple prolines, Cys bridges, or hard-to-synthesize motifs) A composite score was defined to assess peptide favourability: Composite Score = √(Zcharge)2 + (Zhydrophobicity)2 + (ZpI)2, where Z-values represent normalized deviations from ideal ranges (charge: ~0, pI: ~7, hydrophobicity: –0.5 to 0.5). Peptides with low composite scores were considered balanced and favourable.

### 4. 3D Scatter Plot and Heatmap Visualization

Filtered peptides (~5000) were visualized using a 3D scatter plot in OriginPro (X = net charge, Y = hydrophobicity, Z = pI). Color-coding was applied based on the composite score index (bins 1–10), allowing quick visual assessment of physicochemical clustering. Additional heat maps were generated by binning charge and hydrophobicity values against composite scores, enabling 2D insight into peptide distribution trends.

### 5. Peptide Structure Prediction

Top candidates from each bin were selected (n = 100) for structure prediction using AlphaFold2 (via ColabFold) for monomeric peptide folding (28). Predicted structures were energy-minimized and validated using MolProbity (29) and Ramachandran analysis to ensure backbone geometry and stability.

### 6. Protein–Peptide Docking

Molecular docking was performed using MEGADOCK (30) (in the BoltPro platform). Each peptide was docked individually into the active site or extracellular region of its corresponding target. Grid boxes were centered on known ligand-binding residues, and flexible side-chain docking was enabled. Top docking poses were selected based on binding affinity (kcal/mol) and interaction profile (hydrogen bonds, π-π stacking, salt bridges). For cross-target docking, top peptides from one receptor (e.g., TGFBR2) were tested against the remaining targets to identify multifunctional or pan-active peptides.

### 7. Interaction Analysis and Binding Affinity Calculation

Binding interfaces were analyzed using: PyMOL (31) (visualization and contact mapping); PRODIGY (32) (ΔG estimation and dissociation constant); Peptide-protein interaction + (2D interaction diagrams). Key interaction residues were mapped, and contacts with pharmacologically relevant domains (e.g., MMP-1 catalytic zinc-coordination site) were prioritized. Peptides with binding energies < –25 kcal/mol and multi-residue interactions were flagged as high-confidence leads.

## Results

Upon performing sequence-based peptide generation against specific protein targets/ receptors, 5000 different peptide sequences were obtained that were up to 12 residues in length. Property prediction of these generated peptide sequences performed with PepFuNN (27) yielded in the calculation of the net charge, average hydrophobicity, isoelectric point, solubility rules, and synthesis rules.

**Fig 1:**
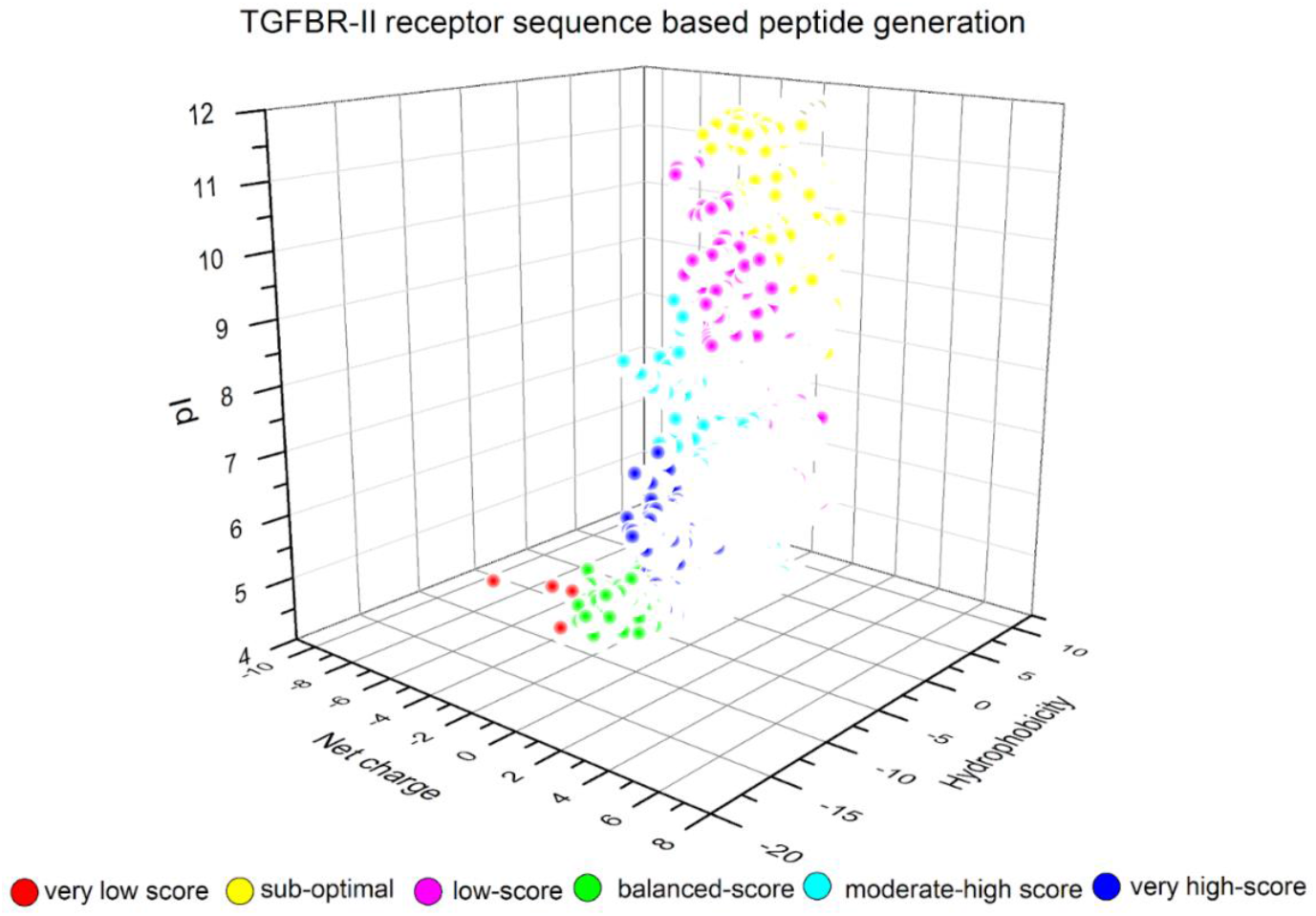
Peptide sequences generated for the TGFBR-II protein (receptor).

The peptide properties generated are represented in a 3-D scatter plot. According to the three significant properties of charge, hydrophobicity, and pI, the peptides were scored, and color bins were assigned to them in the following way:

This way, the gradient helps visually map peptide “favourability” or composite strength. Low score refers to high charge, extreme pI, and very hydrophobic/ hydrophilic. Mid refers to moderate charge, balanced hydrophobicity, pI ~6–8, and high refers to Net charge ~0–±1, mild hydrophobicity, pI ~7, balanced properties for solubility and stability.

**Fig 2:**
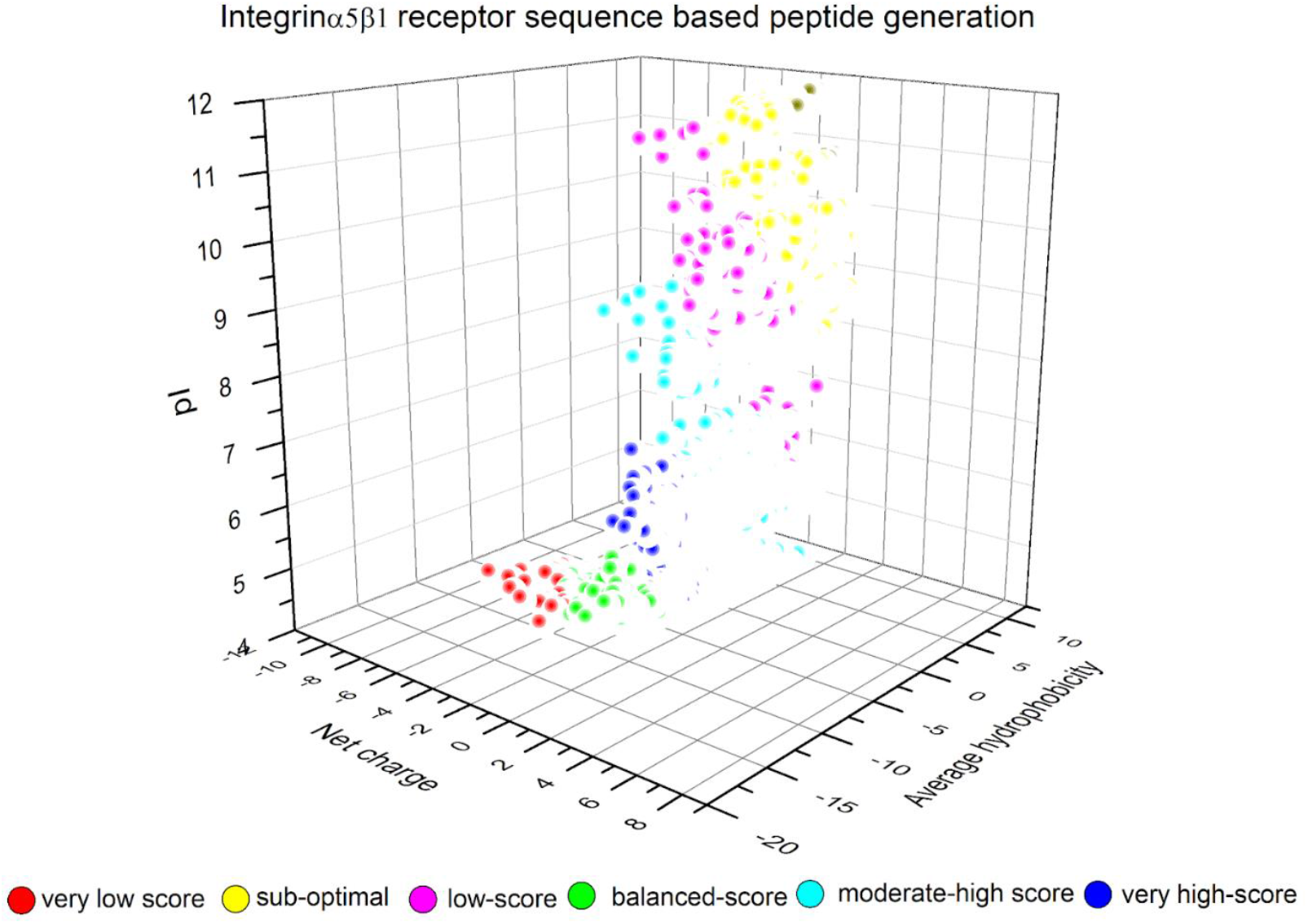
Peptide sequences generated for the Integrinα5β1 protein (receptor).

High net charge, either positive or negative, can enhance binding via electrostatic interactions but may reduce membrane permeability or increase aggregation. Low net charge is generally preferred because it is more neutral and better for solubility and bioavailability.

A peptide with extreme net charge (e.g., +5 or −5) might *lower* the composite score if neutrality is favoured. A peptide with moderate or near-neutral charge might *increase* the composite score if electrostatic balance is desirable.

In the case of average hydrophobicity, while high hydrophobicity can help in membrane penetration or binding to hydrophobic pockets, it often lowers solubility. Similarly, low hydrophobicity improves solubility but may reduce target binding to hydrophobic domains. Peptides with moderate hydrophobicity score higher compared to peptides that are too hydrophobic/ hydrophilic.

Peptides may be less charged in plasma, more stable, and better absorbed when their isoelectric point is close to the physiological pH of ~7.4. In case the pI is very low or very high, the peptide is likely to be highly charged at physiological pH, possibly less stable or prone to aggregation. A composite score favoring plasma stability might rank peptides with pI ≈ 6–8 higher.

**Fig 3:**
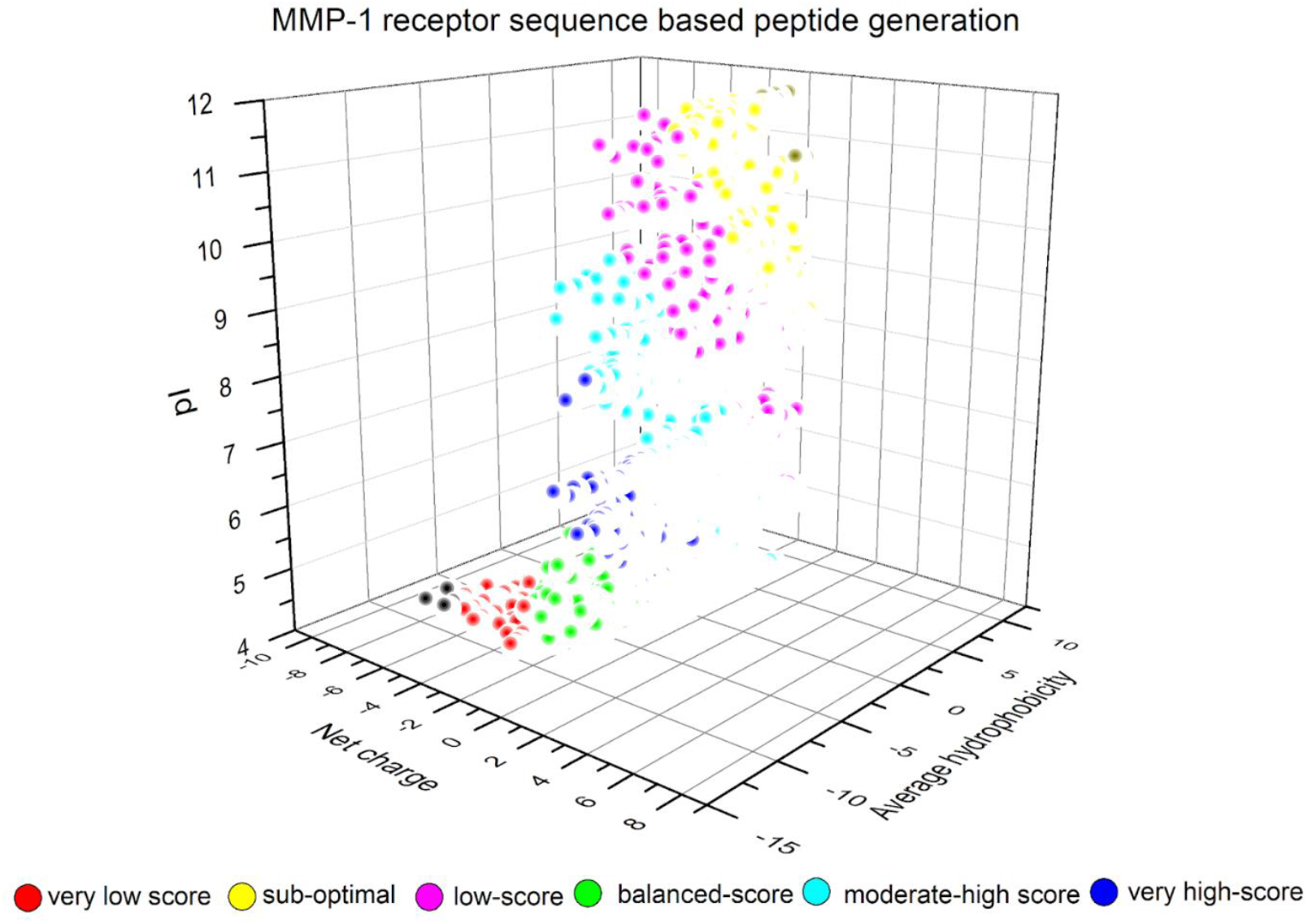
Peptide sequences generated for the MMP-1 protein (receptor).

Some of the common trends observed in the 3D-scatter plots for the three peptide sets of the respective receptors suggested that most data points clustered in the centre of the hydrophobicity and net charge axes, suggesting the moderate hydrophobicity and neutral nature of the peptides. Several yellow and magenta points in the plot can refer to mid-low or slightly low range, suggesting improvement in these peptides. Blue and green dots (higher composite scores) are concentrated around moderate hydrophobicity and net charge ~0 to −2, indicating a sweet spot for balanced peptides. Cyan points are spread out, suggesting they have moderate potential but might vary slightly more across pI or hydrophobicity. Therefore, the green and blue points denote the best peptides, with optimal balance in charge, pI, and hydrophobicity. The 5000 peptides from each set were filtered based on the following characteristics to give rise to the top 10 best-suited peptide candidates selected for structure prediction. A refined filtration criterion was applied to obtain the top 10 peptides that satisfied the following: (i) Net Charge • Mildly positive (0 to +2): Enhances cell penetration without triggering toxicity or instability. (ii)Average Hydrophobicity • 0.5 to 2.0: Avoids excessive hydrophobicity (which could cause aggregation or skin irritation) while maintaining receptor interaction. (iii)Isoelectric Point (pI) • 6.5 to 8.5: Close to physiological pH to enhance topical stability and receptor binding efficacy. (iv) Solubility Rules Failed • ≤1: Better aqueous solubility, which is essential for formulation in cosmetic creams/serums (v) Synthesis Rules Failed • ≤1: Ensures cost-effective and feasible peptide synthesis.

Binding free energy of peptide-protein complexes:

Each receptor’s best-performing peptides demonstrated high-affinity binding within functionally relevant domains:

- TMWDHGFARAL, designed against MMP-1, exhibited the strongest target-specific binding with a docking score of –31.62 kcal/mol, forming interactions near the catalytic zinc-binding cleft involving residues such as His218 and Glu219.
- VWGDQWHYKVW, targeted for TGFBR2, bound the ectodomain interface with a docking score of –19.87 kcal/mol, engaging key hydrogen bonding and hydrophobic contacts.
- SFKTVLLDVGK, designed for Integrin α5β1, displayed the highest overall docking affinity of –34.5 kcal/mol, interacting with residues in the beta-propeller domain involved in extracellular ligand recognition.

Peptide generated against TGFBR-II receptor VWGDQWHYKVW: Docked against TGFBR-II, Integrin α5β1, MMP-1

| **TGFBR-II** | **Integrin $\alpha 5\beta 1$** | **MMP-1** |
| --- | --- | --- |
| -19.87 kcal/mol | -23.62 kcal/mol | -32.07 kcal/mol |

Peptide generated against Integrin α5β1 receptor SFKTVLLDVGK: Docked against Integrin α5β1, TGFBR-II, MMP-1

| **Integrin $\alpha 5\beta 1$** | **TGFBR-II** | **MMP-1** |
| --- | --- | --- |
| -34.5 kcal/mol | -30.88 kcal/mol | -30.92 kcal/mol |

Peptide generated against MMP-1 receptor TMWDHGFARAL: Docked against MMP-1, TGFBR-II, Integrin α5β1

Peptide generated against MMP-1 receptor TMWDHGFARAL :Docked against MMP-1, TGFBR-II, Integrin $\alpha 5\beta 1$
| **MMP-1** | **TGFBR-II** | **Integrin $\alpha 5\beta 1$** |
| --- | --- | --- |
| -31.62 kcal/mol | -20.15 kcal/mol | -29.44 kcal/mol |

### Cross-Target Binding Evaluation

To assess multifunctionality, the top-scoring peptides were cross-docked against all three receptors. Notably, SFKTVLLDVGK retained strong binding across all three targets (–34.5, – 30.88, and –30.92 kcal/mol for Integrin, TGFBR2, and MMP-1, respectively), indicating pan-target potential. VWGDQWHYKVW showed enhanced affinity toward MMP-1 (–32.07 kcal/mol) and Integrin α5β1 (–23.62 kcal/mol), despite being designed for TGFBR2, suggesting structural compatibility beyond its original target. Other peptides, such as LDCLGLHMRWQ and LEHSITGHKYI, also showed cross-target affinities exceeding –25 kcal/mol, positioning them as potential dual-function candidates.

### Interaction Profiling

Detailed interaction maps confirmed that top peptides engaged residues within receptor epitope regions and catalytic sites, especially in MMP-1 and Integrin. Hydrogen bonding, electrostatic contacts, and π–π stacking interactions were commonly observed, particularly involving charged and aromatic residues in the peptides (e.g., Lys, Arg, Trp, Tyr).

Peptide generated against TGFBR-II receptor VWGDQWHYKVW docked against TGFBR-II, VWGDQWHYKVW engages TGFBR-II through aromatic stacking (Phe255), moderate hydrophobic burial, and water-mediated polar contacts. The combined contacts suggest surface groove engagement, possibly in the receptor’s ectodomain or dimerization interface. These interactions are non-catalytic but stabilizing, ideal for modulation rather than inhibition, consistent with a cosmeceutical or signaling-regulatory role.

**Fig 4:**
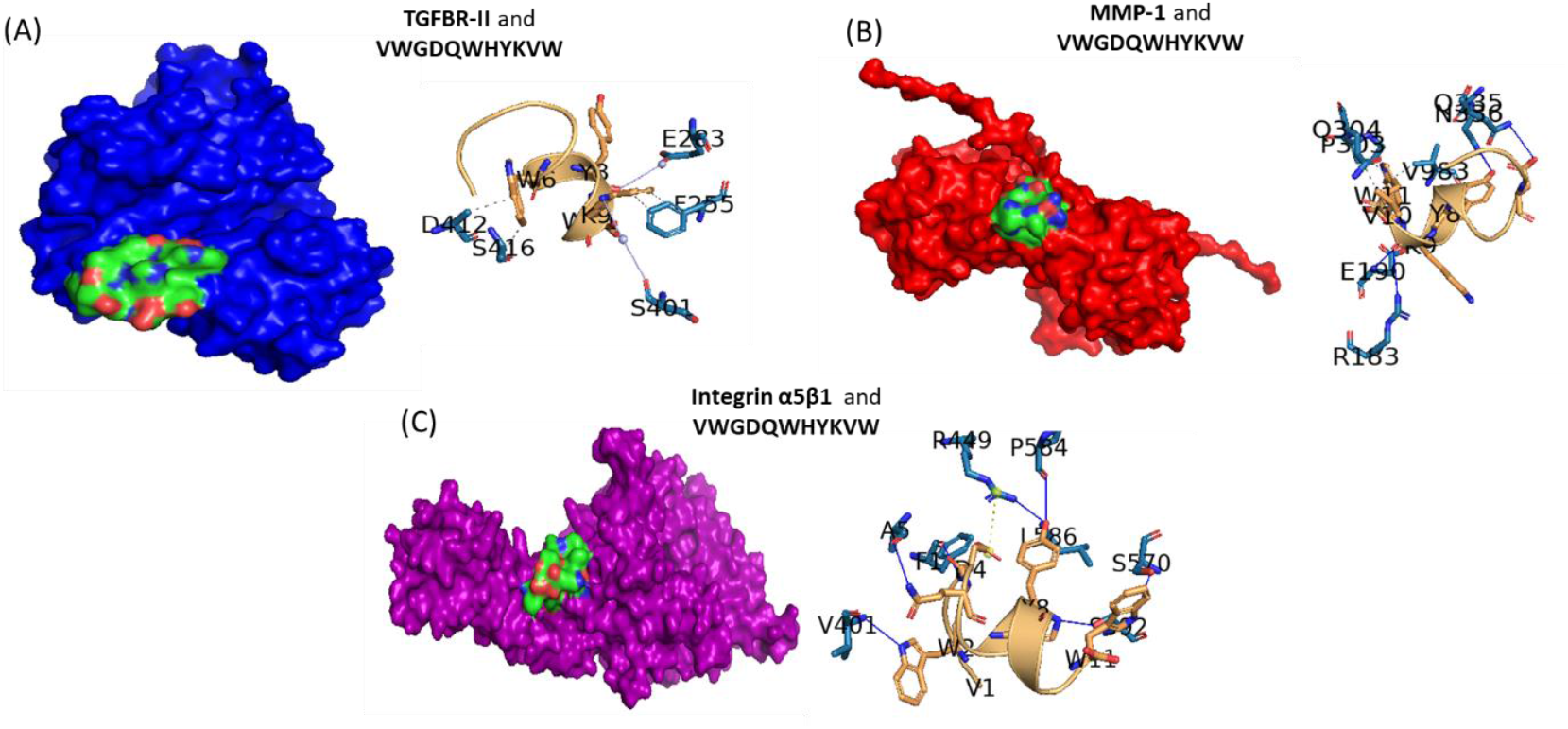
Docking of peptide TMWDHGFARAL to receptors (A) TGBR-II, (B) MMP-1, and (C) Integrin α5β1. The surface representation for each receptor-peptide (protein-ligand) complex and the respective interacting residues have been depicted. TGFBR-II is color-coded blue, MMP-1 is color-coded red, and Integrin α5β1 is color-coded purple in the surface representations. In the protein-peptide residue interaction profile, the protein residues interacting are shown as blue sticks, and the peptide residues are in light brown.

Four key hydrophobic interactions were observed, stabilizing the TGFBR-II– VWGDQWHYKVW complex. The Phe255 (3.62 Å, 3.63 Å) has a dual contact nature of interaction, likely engaging in π–π stacking or CH–π interactions with Tyr or Trp in the peptide. The Asp412 (3.49 Å) interaction may involve side-chain aliphatic interactions with non-polar residues in Valine, and the Ser416 (3.74 Å) is likely a weak interaction with the non-polar surface of the peptide or the backbone atoms. The peptide is anchored at the surface groove of TGFBR2 through hydrophobic interactions involving Phe255, suggesting engagement with aromatic side chains (like Trp or Tyr) from the peptide. These interactions help stabilize surface-level docking in the ectodomain. Two water-mediated bridges contribute to complex stability, bridging the ligand and protein via solvent molecules. The Glu238 residue of the receptor TGFBR-II is the hydrogen bond donor in the water bridge, in a case of classic bridging. That is, the protein donates a hydrogen to the water molecule, which then interacts with the ligand (peptide) acceptor, with the acceptor to water distance of 3.63 Å, and donor to water distance of 2.84 Å, and an overall donor angle of 135.24**°**. In another case of reverse bridging, residue Ser401 of the receptor forms a peptide-to-protein water bridge via the N-terminal group, stabilising the peptide orientation, with acceptor to water distance of 3.42 Å, and donor to water distance of 3.46 Å, and an overall donor angle of 141.70**°**. Therefore, in this case, the water molecule accepts a hydrogen from the ligand and then donates one to the protein acceptor. The presence of two water bridges adds flexibility and entropic favourability, with one Glu-mediated and one peptide-driven bridge. These contacts complement the hydrophobic interactions, enabling a balanced binding mode.

Eight hydrogen bonds were detected for the Integrin α5β1-VWGDQWHYKVW complex, as shown in Table 5. Bonds under 3.0 Å (especially < 2.5 Å) are strong — e.g., those from Phe1, Ala5, Arg449. ARG 449A plays a key dual role of forming a hydrogen bond and participating in salt bridge formation. While, interactions 3, 4, 5, and 8 involve the receptor/ protein Integrin α5β1 acting as the donor, in interactions 1, 2, 6, and 7, the ligand (peptide) is donating the hydrogen instead; the protein is the acceptor. Protein residue hydrogen bond donor indicates that the protein forms a stable anchoring interaction with the peptide. The ligand residues acting as the hydrogen bond donor may indicate the peptide is initiating the contact, possibly to stabilize its orientation in the binding pocket.

**Table 1:** Scoring and color scheme of all generated peptide sequences for 3 different proteins.

| Bin | Score Range | Suggested Color |
| --- | --- | --- |
| 1 | Lowest | Red |
| 2–4 | Low-Mid | Orange |
| 5–7 | Mid | Yellow |
| 8–9 | High | Cyan–Green |
| 10 | Highest | Blue |

**Table 2:** Legend for composite scoring.

| Composite Score Range | Color | Interpretation |
| --- | --- | --- |
| < 2 | Red | Very low: Highly charged or extreme pI/hydrophobicity (possibly unstable) |
| ~2.75–4.25 | Magenta | Slightly low score (mild imbalance) |
| ~3.5–4.25 | Yellow | Low: moderately sub-optimal |
| ~4.25–5 | Green | Moderate composite score (balanced properties) |
| ~5–5.75 | Cyan | Moderate-high score |
| ~6.5–7.25 | Blue | High score (favorable net charge/pI/hydrophobicity) |

**Table 3:**
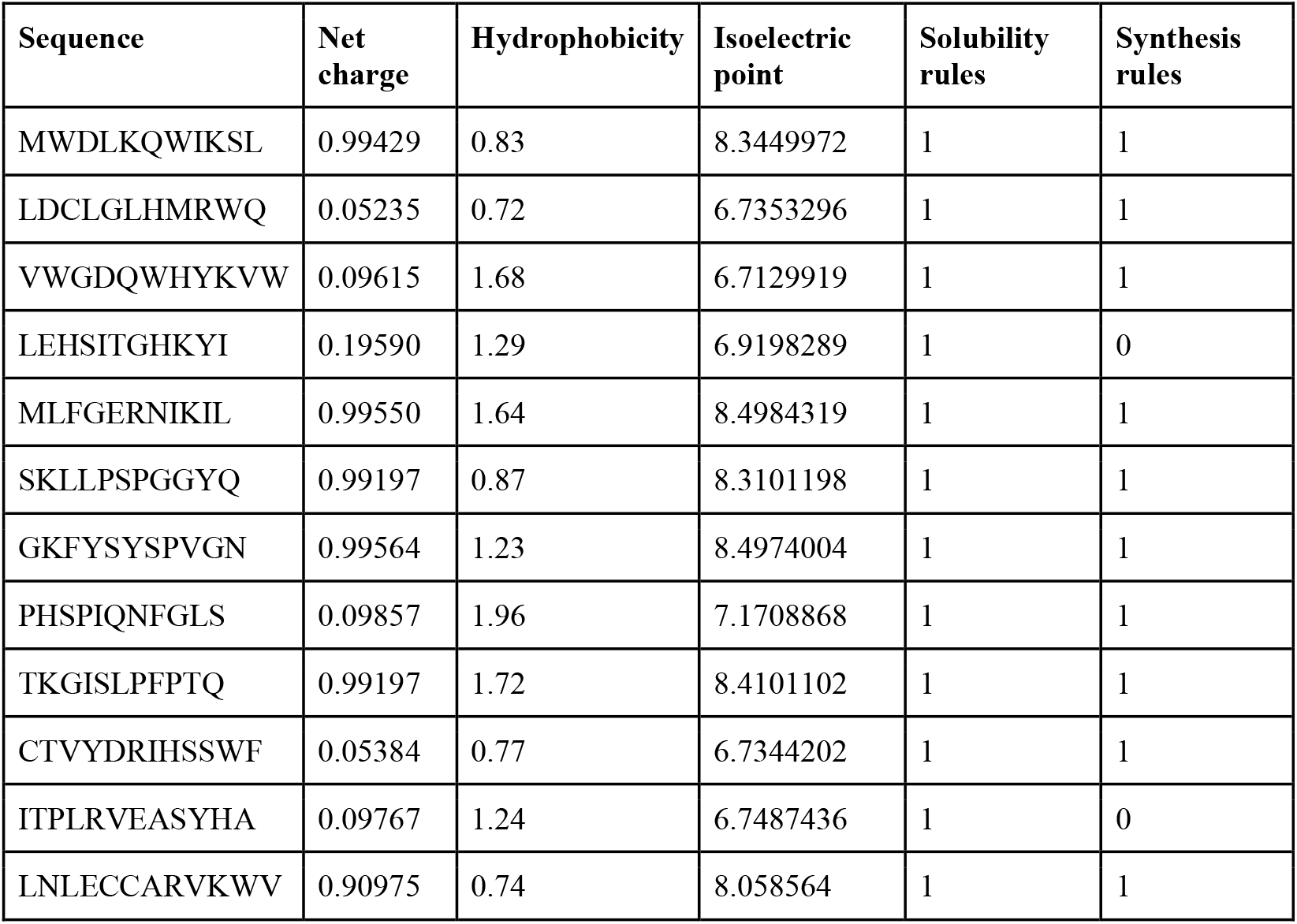
TGBFR-II receptor-based peptides generated and filtered.

**Table 3:**
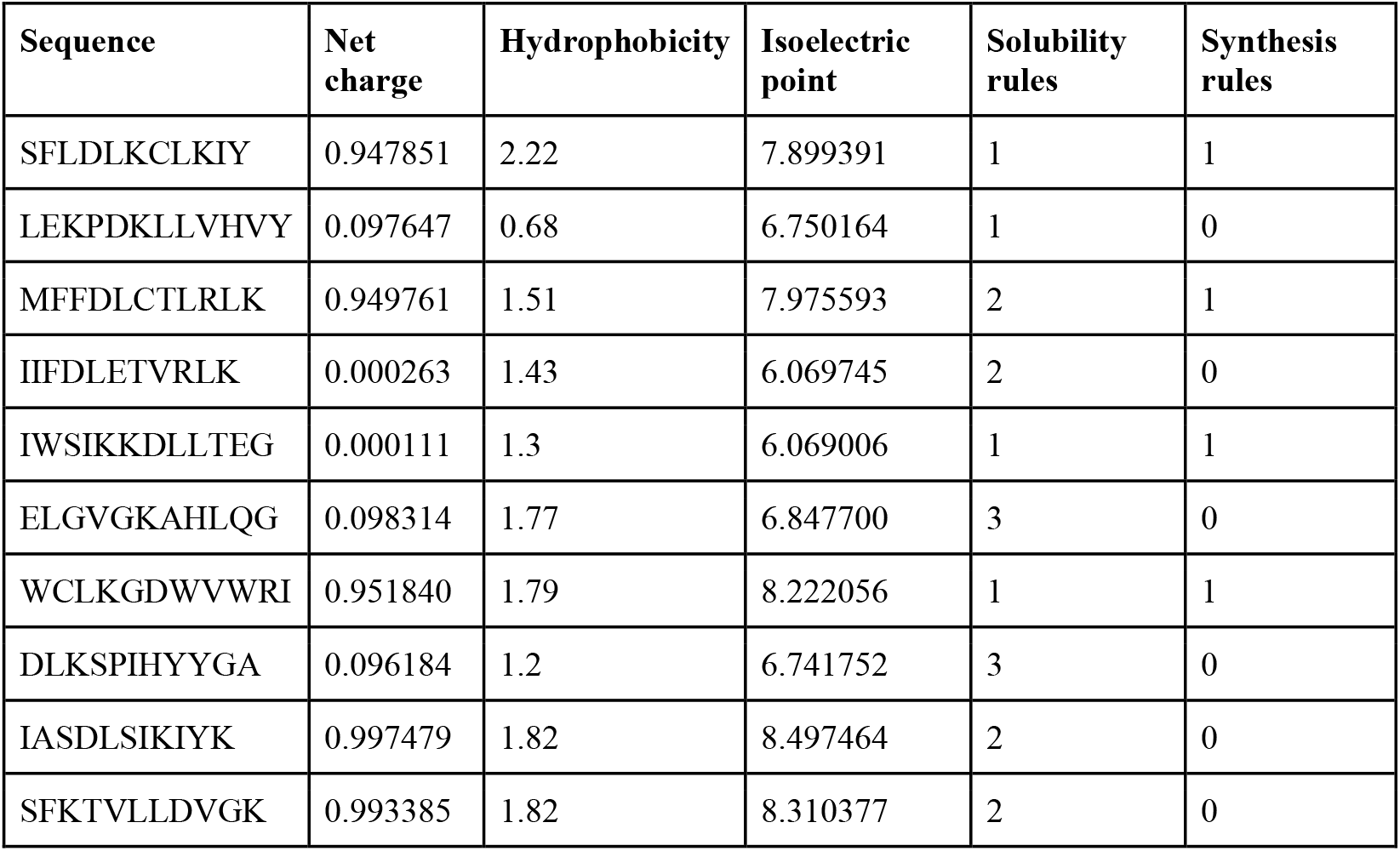
Integrin α5β1 receptor-based peptides generated and filtered.

**Table 4:**
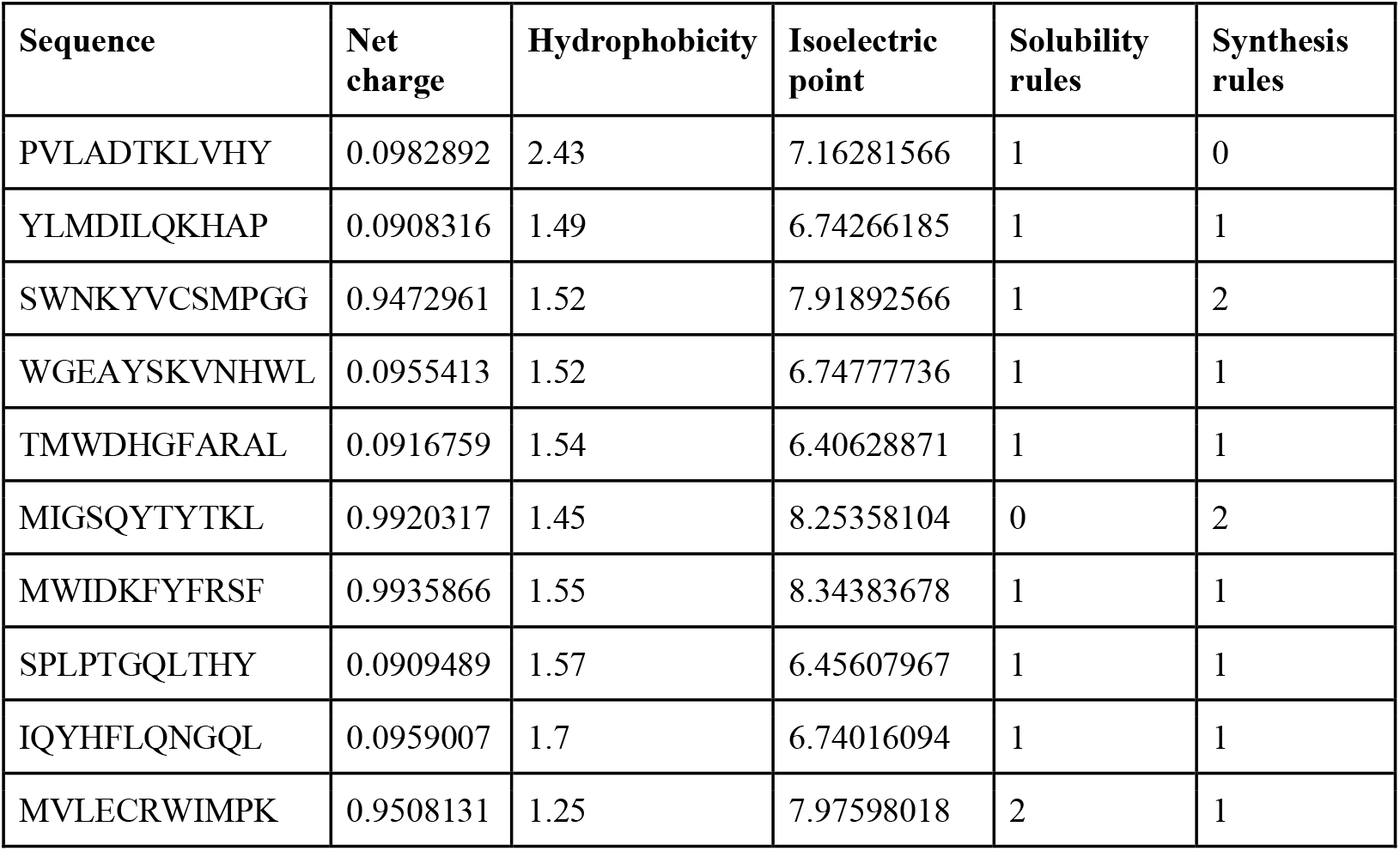
MMP-1 receptor based filtered peptides.

**Table 5:** There are 8 hydrogen bonds between VWGDQWHYKVW and the Integrin protein.

| Residue (Chain ID) | AA | Distance H-A (Å) | Protein Donor? | Side Chain? |
| --- | --- | --- | --- | --- |
| 1A | PHE | 2.04 | × | × |
| 5A | ALA | 2.50 | × | × |
| 401A | VAL | 2.95 | ✓ | × |
| 449A | ARG | 2.64 | ✓ | ✓ |
| 570A | SER | 2.87 | ✓ | ✓ |
| 584A | PRO | 2.84 | × | × |
| 586A | LEU | 2.86 | × | × |
| 592A | SER | 3.27 | ✓ | ✓ |

Only one salt-bridge is detected, where Arg449 of the Integrin α5β1 interacts with Asp4 group on the peptide (VWGDQWHYKVW), forming a salt bridge, typically a strong electrostatic interaction. The 5.38 Å distance suggests it’s likely weak-to-moderate in strength (ideal < 4 Å). This residue is a hotspot, stabilizing the complex via both H-bond and electrostatic interaction. The peptide forms strong and diverse interactions with integrin α5β1, notably through ARG 449A, SER 570A, and VAL 401A. These interactions suggest good binding affinity and specificity, especially with dual-interacting residues like ARG 449A.

There are three hydrophobic interactions between the MMP-1 protein/ receptor and the peptide generated against TGFBR-II receptor VWGDQWHYKVW. Pro303 (3.15 Å), Gln304 (3.31 Å), and Val983 (3.29 Å) are involved in the hydrophobic interactions that support van der Waals stability. While proline and Valine are classic hydrophobic interactors, all the interaction distances fall within an ideal range of 3.1–3.3 Å, which is suitable for effective packing without steric clash.

Also, five hydrogen bonds (Table 6) are identified between the MMP-1 protein/ receptor and the peptide VWGDQWHYKVW. Gln335 is particularly important: It participates in two H-bonds, including a very strong one at 1.83 Å with near-linear geometry (176.86°). Arg183 and Glu190 form stable hydrogen bonds via charged groups, supporting electrostatic stabilization. Most donor atoms are from protein, showing that MMP-1 is acting primarily as a hydrogen bond donor. The strongest hydrogen bond is Gln335A (1.83 Å, 176.86°), which indicates a near-ideal H-bond geometry. Gln335 and Asn336 are interaction hotspots, with multiple or high-quality interactions. The peptide binding pocket in MMP-1 exhibits both hydrophobic and polar characteristics, allowing for a complementary fit.

**Table 6:** 5 hydrogen bonds between the VWGDQWHYKVW and MMP-1 protein exist.

| Residue (Chain ID) | AA | Distance H-A (Å) | Donor Angle | Protein Donor? | Side Chain? |
| --- | --- | --- | --- | --- | --- |
| 183A | ARG | 3.32 | 122.89 | ✓ | ✓ |
| 190A | GLU | 2.09 | 118.24 | ✓ | ✗ |
| 335A | GLN | 1.83 | 176.86 | ✓ | ✓ |
| 335A | GLN | 3.24 | 107.62 | ✗ | ✗ |
| 336A | ASN | 3.03 | 146.48 | ✓ | ✗ |

**Table 7:** Five hydrophobic contacts exist between SFKTVLLDVGK and Integrin α5β1.

| Residue (Chain ID) | AA | Distance (Å) | Ligand Atom | Protein Atom |
| --- | --- | --- | --- | --- |
| 164F | LEU | 3.99 | 7905 | 7480 |
| 164F | LEU | 3.36 | 7936 | 7479 |
| 189F | PRO | 3.25 | 7926 | 7661 |
| 189F | PRO | 3.29 | 7937 | 7661 |
| 192F | THR | 3.94 | 7903 | 7680 |

**Table 8:**
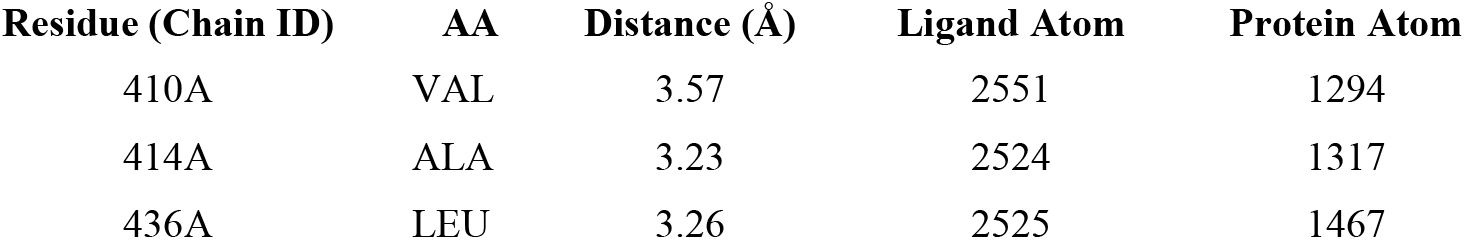
Three hydrophobic contacts exist between SFKTVLLDVGK and TGFBR-II.

**Table 9:** Two hydrogen bonding interactions between SFKTVLLDVGK and TGFBR-II.

| Residue (Chain ID) | AA | Distance H-A (Å) | Distance D-A (Å) | Donor Angle (°) | Protein Donor? | Side Chain? |
| --- | --- | --- | --- | --- | --- | --- |
| 414A | ALA | 3.32 | 4.07 | 131.63 | ✗ | ✓ |
| 432A | SER | 2.87 | 3.70 | 143.24 | ✓ | ✓ |

**Table 10:** Five hydrophobic contacts between SFKTVLLDVGK-MMP-1.

| Residue (Chain ID) | AA | Distance (Å) | Ligand Atom | Protein Atom |
| --- | --- | --- | --- | --- |
| 188A | PHE | 3.87 | 3273 | 852 |
| 188A | PHE | 3.69 | 3272 | 854 |
| 190A | GLU | 3.72 | 3303 | 874 |
| 303A | PRO | 3.47 | 3270 | 1774 |
| 982C | ILE | 2.47 | 3280 | 3127 |

**Table 10:** Five hydrophobic contacts between SFKTVLLDVGK-MMP-1.

| Residue (Chain ID) | AA | Distance H-A (Å) | Distance D-A (Å) | Donor Angle (°) | Protein Donor? | Side Chain? |
| --- | --- | --- | --- | --- | --- | --- |
| 335A | GLN | 3.56 | 4.05 | 113.54 | ✓ | ✓ |
| 336A | ASN | 2.57 | 3.49 | 155.60 | ✓ | ✗ |

**Table 11:** Five hydrogen bonding interactions between TMWDHGFARAL-MMP-1.

| Residue | AA | Distance H-A (Å) | Distance D-A (Å) | Donor Angle | Protein Donor? | Side Chain? |
| --- | --- | --- | --- | --- | --- | --- |
| 287A | ASN | 3.42 | 4.04 | 123.11 | ✓ | ✓ |
| 314A | GLU | 2.67 | 2.99 | 100.07 | ✗ | ✗ |
| 316A | ALA | 3.05 | 3.99 | 161.55 | ✓ | ✓ |
| 364A | GLU | 1.40 | 2.19 | 128.69 | ✓ | ✗ |
| 365A | GLU | 3.42 | 3.78 | 103.67 | ✓ | ✗ |

**Table 12:** 3 hydrogen bonds detected between the TMWDHGFARAL-TGFBR-II complex.

| Residue | AA | Distance H-A (Å) | Distance D-A (Å) | Donor Angle | Protein Donor? | Side Chain? |
| --- | --- | --- | --- | --- | --- | --- |
| 432A | SER | 2.50 | 3.39 | 152.00 | ✗ | ✓ |
| 435A | ASN | 3.34 | 3.66 | 100.63 | ✓ | ✓ |
| 436A | LEU | 3.33 | 3.92 | 120.19 | ✓ | ✗ |

Overall, the interaction between MMP-1 and VWGDQWHYKVW is a high-affinity interaction, driven by well-oriented hydrogen bonds and close hydrophobic contacts. Residues like Gln335, Arg 183, and Pro303 are likely critical for binding specificity and can be considered for future mutational analysis or peptide optimization.

**Fig 5:**
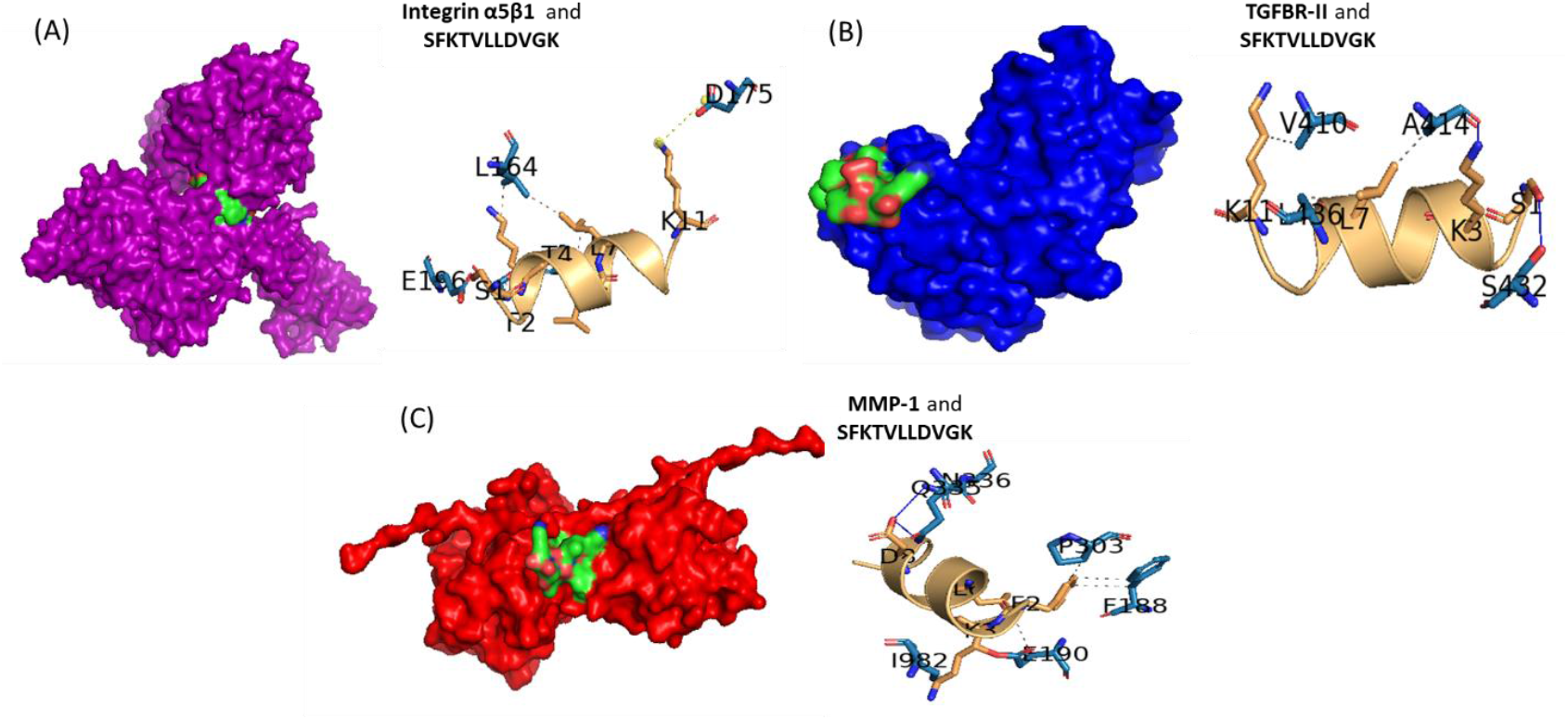
Docking of peptide SFKTVLLDVGK to receptors (A) Integrin α5β1, (B)TGFBR-II, and (C) MMP-1. The surface representation for each receptor-peptide (protein-ligand) complex and the respective interacting residues have been depicted. Integrin α5β1 is color-coded purple, TGFBR-II is color-coded blue, and MMP-1 is color-coded red in the surface representations. In the protein-peptide residue interaction profile, the protein residues interacting are shown as blue sticks, and the peptide residues are shown in light brown color.

A total of 5 hydrophobic contacts were observed, with key involvement from Leu164 and Pro189 residues in the Integrin α5β1 protein and SFKTVLLDVGK complex. Pro189 makes two hydrophobic contacts, suggesting its pivotal role in the peptide’s anchoring. Leu164 also makes dual interactions, which are typical for stabilizing hydrophobic packing. Hydrophobic contacts range from 3.25–3.99 Å, all within the ideal non-bonded interaction distance. Only one hydrogen bond was detected in the complex, with relatively weak geometry (long distance and lower angle) at Glu196 with hydrogen to acceptor distance 3.65 Å, and donor to acceptor distance of 4.02 Å and angle 105.52°. Only one salt bridge was observed between Asp175 of the protein and Lys11 of the peptide. With a distance of 5.09 Å, this salt bridge is borderline in strength (ideal is < 4 Å), but still contributes to electrostatic stabilization. This implies the peptide relies more on hydrophobic stabilization than polar or electrostatic bonding. The protein is negatively charged here (ASP), while the ligand group (Lysine) is positively charged, supporting a typical charge complementarity. Hydrophobic interactions dominate the binding landscape, with dual involvement from Leu164 and Pro189 residues, suggesting this peptide may anchor through hydrophobic embedding rather than extensive polar interactions. The complex may benefit from optimization of polar contacts or stronger salt bridge formation (e.g., through side-chain modifications or D-amino acid substitutions).

The peptide SFKTVLLDVGK docked against TGFBR-II displays three hydrophobic contacts, all within favourable interaction distances (3.23–3.57 Å), suggesting optimal packing, contributing favourably to stability. All residues involved are small-to-medium non-polar amino acids (Val410, Ala414, Leu436), consistent with van der Waals-driven binding. 2 hydrogen bonds at Ser432 forms a solid H-bond (2.87 Å, 143.24°), indicating good orientation and strength, and at Ala414 that forms a weaker bond (longer distance, protein not donating), possibly electrostatically driven.

The peptide SFKTVLLDVGK binds modestly well to TGFBR-II, mainly driven by Hydrophobic contacts with small aliphatic residues, and a single well-aligned hydrogen bond from SER 432A. The interaction profile suggests lower polar stabilization than seen in previous targets like Integrin or MMP-1. This introduces potential enhancement via sequence edits (e.g., introducing polar residues at the binding interface).

The SFKTVLLDVGK-MMP-1 complex showcases five hydrophobic contacts and two hydrogen bonding contacts. Phe188 forms two contacts (π-π or hydrophobic stacking), and Ile982 has the shortest interaction distance (2.47 Å), suggesting a very tight fit in the binding pocket. Glu190, which is typically charged, appears here as a non-polar face participant, indicating a partial hydrophobic surface involved in interaction. Among the hydrogen bonds, Asn336 forms a strong and linear hydrogen bond (155.6° and 2.57 Å), a significant stabilizer, while Gln335 contributes a weaker but still stabilizing H-bond. Both donors are from the protein backbone or side chain, not the ligand.

The peptide SFKTVLLDVGK shows a robust binding profile with MMP-1, supported by tight hydrophobic packing (Ile982, Pro303, Phe188), and strong hydrogen bonding, especially via Asn336. This interaction pattern indicates better affinity and stability between the SFKTVLLDVGK-MMP-1 complex compared to its binding with TGFBR**-**II, where hydrogen bonding was weaker.

The peptide generated against the MMP-1 receptor is the peptide TMWDHGFARAL, and when docked against MMP-1, forms four hydrophobic contacts, spanning aromatic, polar, and aliphatic residues. Two Phe residues (289, 315) anchor the peptide via strong aromatic hydrophobic interactions. Surprisingly, Glu 365 and ASP 414 (typically polar/charged) contribute hydrophobically, likely via non-polar side chain surfaces or partial burial.

**Fig 3:**
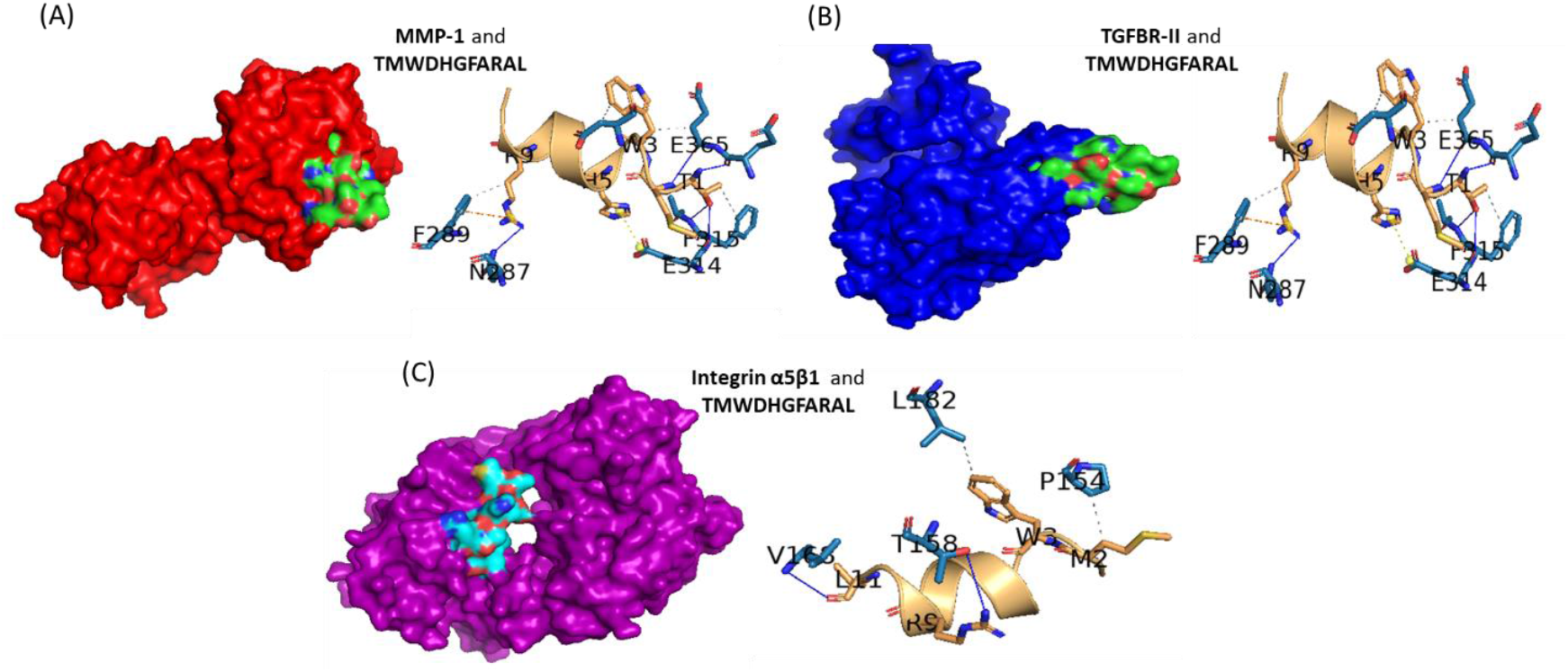
Docking of peptide TMWDHGFARAL to receptors (A) MMP-1, (B) TGBR-II, and (C) Integrin α5β1. The surface representation for each receptor-peptide (protein-ligand) complex has been depicted along with the respective interacting residues. MMP-1 is color coded red, TGFBR-II is color coded blue, and Integrin α5β1 is color coded purple in the surface representations. In the protein-peptide residue interaction profile, the protein residues interacting are shown as blue sticks, and the peptide residues are shown in light brown color.

Also, five hydrogen bonds are detected for the MMP-1-TMWDHGFARAL complex, where Glu364A forms a very strong H-bond (1.40 Å), a central stabilizer. Also, Ala316 shows an almost linear geometry (161.55°), which is also favourable. The donor atoms are essentially from protein side chains or backbone, emphasizing protein-driven stabilization. Additionally, there is one π–cation interaction between Phe 289 of the MMP-1 protein and the Arg9 of the peptide, indicative of electrostatic and aromatic stabilization, enhancing binding specificity. Salt bridge formed between the Glu314 of the MMP-1 protein and the His5 of the TMWDHGFARAL protein adds electrostatic strength, although 4.44 Å is at the upper range for strong ionic interaction. TMWDHGFARAL exhibits a highly favourable interaction profile with MMP-1, likely leading to strong binding affinity.

The combination of diverse stabilizing forces—π–cation, salt bridge, H-bonds—alongside aromatic and aliphatic packing makes this peptide, TMWDHGFARAL, a strong candidate for MMP-1 inhibition or targeting.

The peptide TMWDHGFARAL was also docked against the TGFBR-II receptor to analyse the binding affinity and interaction. It was observed that only one hydrophobic interaction was formed between Asp411 of the protein and the peptide. Asp typically forms polar or ionic interactions, but here it’s involved in a weak hydrophobic contact, indicating a less pronounced non-polar pocket compared to MMP-1. Three hydrogen bonds belonging to Ser432, which provides the strongest and most linear H-bond (2.50 Å, 152°), Leu436, and Asn435 that also contribute, though with more moderate geometry, were detected. TGFBR-II contributes two donor residues (Asn435, Leu436), stabilizing peptide binding moderately. The two water bridges formed serve as extended H-bonds, improving stability in aqueous environments. Met434 forms an almost ideal water bridge (tight angles, short distances), and the Glu437 bridge is moderately favourable.

The peptide TMWDHGFARAL binds modestly to TGFBR-II, with hydrogen bonds and water bridges compensating for limited hydrophobic anchoring. The best stabilizing features are the direct H-bond from Ser432 and the water-mediated bridge via Met434. Compared to the stronger and multi-modal binding of TMWDHGFARAL with MMP-1, this interaction with TGFBR-II is weaker and more polar-dependent.

The complex between the peptide generated against the MMP-1 receptor TMWDHGFARAL and the protein Integrin α5β1 showcased two hydrophobic interactions. Pro154 and Leu182, both aliphatic, contribute van der Waals interactions with distances 3.57–3.76 Å, which are within acceptable hydrophobic contact range, though not as tight as an ideal contact distance (<3.5 Å). These interactions are therefore consistent with moderate hydrophobic packing. Two hydrogen bonds, Val168 and Thr158, were also detected in the TMWDHGFARAL-Integrin α5β1 complex. Val168 provides a relatively strong hydrogen bond with decent geometry (140.34°). Thr158 contributes a weaker H-bond, likely stabilizing via side-chain hydroxyl interaction. The hydrogen bonding is limited in number and strength compared to TGFBR-II and MMP-1 interactions, and the peptide TMWDHGFARAL shows weaker interaction strength with Integrin α5β1 compared to MMP-1 and TGFBR-II. This complex also lacks strong hydrophobic anchoring and high-quality polar contacts. While binding may still be biologically relevant, it could benefit from sequence optimization to improve integrin affinity.

Unlike many conventional approaches focusing on single-target peptide design, this work integrates multi-target screening, physicochemical filtering, and docking-based structural validation to identify peptides with pan-target efficacy.

A critical strength of the study lies in the multi-parametric composite scoring system, which accounts for charge, hydrophobicity, pI, solubility, and synthetic feasibility—properties that directly affect formulation stability, skin permeability, and bioavailability in topical applications. The 3D composite visualization further enabled rational clustering and selection of peptide candidates that lie within a ‘sweet spot’ of physicochemical balance.

Structural predictions using AlphaFold2 and PEP-FOLD3, followed by protein–peptide docking via HADDOCK and AutoDock Vina, allowed for mechanistic insight into interaction specificity. Binding free energies across all peptides ranged from –19.8 to –34.5 kcal/mol, with several peptides showing robust cross-target binding.

Among the top performers:

- SFKTVLLDVGK, designed for Integrin α5β1, showed pan-target potential, binding all three receptors with high affinity (≥ –30 kcal/mol).
- VWGDQWHYKVW, designed for TGFBR-II, also showed strong binding to MMP-1 and Integrin, suggesting a flexible binding motif architecture.
- TMWDHGFARAL, optimized for MMP-1, demonstrated high binding stability via π– cation, salt bridge, and hydrophobic interactions—ideal for protease inhibition.

Interaction profiling further confirmed the engagement of catalytically relevant domains, hydrogen bonding with key side chains, and stability-enhancing interactions such as water bridges. This highlights the dual design utility: target specificity and topical compatibility.

However, it is essential to note that while in silico affinity and interaction patterns are strong indicators, they must be complemented with in vitro bioactivity, cell penetration assays, and degradation kinetics to translate these peptides into practical formulations.

## Conclusions

This work successfully demonstrates a rational, multi-step computational design pipeline for generating short, stable, multifunctional peptides that can act across multiple skin-aging pathways. By integrating de novo peptide generation, machine learning–based filtering, composite property scoring, structural prediction, and docking, the study identified novel peptide candidates capable of stimulating collagen synthesis (via TGFBR-II engagement), enhancing fibroblast-ECM interaction (via Integrin α5β1), and inhibiting matrix degradation (via MMP-1 suppression). Among the candidates, SFKTVLLDVGK and TMWDHGFARAL emerge as lead peptides with broad-spectrum binding and balanced physicochemical properties. This integrative pipeline not only enables peptide optimization for cosmeceutical use but also serves as a template for future multifunctional bioactive design, potentially extendable to wound healing, scar modulation, or even anti-inflammatory applications.

## Notes

### Competing Interest Statement

The authors have declared no competing interest.

